# From blood to pluripotency: Fibrocytes as a reprogrammable somatic cell source for bovine iPSCs

**DOI:** 10.1101/2025.10.30.685680

**Authors:** Hannah Sylvester, Prasanthi P. Koganti, Shailesh Gurung, Viju V. Pillai, Vimal Selvaraj

## Abstract

Fibrocytes represent a distinct somatic cell type derived from peripheral blood leukocytes, first described as spindle-shaped adherent cells with dual hematopoietic and mesenchymal features. Although fibrocytes were identified in early descriptive studies across several mammalian systems, most of this work predated modern molecular approaches, and the cells remain incompletely defined at the molecular level and have not previously been derived or characterized in cattle. Seeking somatic cells that could be collected aseptically and reproducibly under field conditions for reprogramming to pluripotency, we recognized fibrocytes as a practical and previously unexplored candidate population. Here, we establish a reproducible method for fibrocyte derivation and expansion from adult bovine blood and define their molecular identity using transcriptomic and network analyses. Principal component and differential expression analyses revealed extensive immune, inflammatory, metabolic, and stress-responsive pathways that distinguished fibrocytes from fibroblasts. Upstream regulator analysis identified a fibrocyte-restricted transcriptional network governed by SPI1, IRF5/IRF7, NFKBIZ, PRDM1, CIITA, and MAFB, supporting a monocyte-derived origin and indicating some retention of hematopoietic lineage memory despite acquisition of mesenchymal features. Optimized fibrocyte medium (FbC; dexamethasone, ascorbate, PDGF-BB, EGF, A83-01, CHIR99021) supported stable proliferation and selectively enhanced cytoskeletal and matrix-constructive programs while attenuating inflammatory tone. When reprogrammed with polycistronic OCT4-SOX2-KLF4-cMYC and SV40 large T antigen, fibrocytes generated induced pluripotent stem cell (iPSC) colonies exhibiting defining molecular and morphological features of pluripotency. These findings establish fibrocytes as a field-adaptable, stably expandable, and reprogrammable somatic cell type with practical applications in induced pluripotent stem cell generation, genetic preservation, and reproductive biotechnology.

## Introduction

The concept of circulating cells in blood contributing to tissue repair dates back over a century. In 1922, it was demonstrated that cells assuming the appearance of fibroblasts could be cultured from blood of adult chickens [1]. At the time, the origin of these cells remained speculative, with postulates suggesting a transformation of circulating monocytes, or a fibroblastic cell type that exist in small numbers in peripheral blood. In 1927, it was demonstrated that fibroblast-like cells with indistinguishable morphology from connective tissue fibroblasts could be cultured from guinea pig blood [2]. These observations were later extended to human peripheral blood in 1958, confirming that fibroblast-like cells could be propagated from cells in adult human circulation [3]. Supporting these findings, radiolabeled tracer studies linked circulating leukocytes to new connective tissue formation during wound repair [4]. Further validation came from experiments in which human mononuclear leukocytes, cultured in diffusion chambers in vivo, differentiated into “polyblasts” before assuming the morphological characteristics of collagen-producing fibroblast-like cells [5,6]. These results were consistently reproduced across different mammalian species using various methodologies [7,8]. While the precise cellular origin of these fibroblast-like cells remained uncertain, the consensus emerged that a subset, termed “fibrocytes,” originated from circulating mononuclear cells and contributed to wound repair [9].

The search for the precise identity of fibrocytes in peripheral blood remained an active area of investigation for several decades. Although the notion of fibrocytes as a *de novo* rare leukocyte subpopulation remains plausible [10], the extended culture times required for their detection do not exclude the likely possibility of trans-differentiation in their derivation. Subsequent studies have demonstrated the transformation of monocytes into fibroblast-like fibrocytes, solidifying their identity as neo-fibroblasts [11]. Additionally, fibrocytes were found to retain epigenetic markers indicative of their hematopoietic lineage. They continued to express overlapping surface markers such as CD11b, CD13, CD34, and CD45 [10] and exhibited antigen-processing and presenting capabilities similar to monocytes [12]. Despite these similarities, fibrocytes were functionally distinct due to their pronounced role in extracellular matrix deposition and their involvement in harmonizing both inflammatory and reparative phases of wound healing [13]. Further research has elucidated key signaling pathways governing fibrocyte differentiation and migration, shedding light on their recruitment to wound sites and their dynamic role in tissue repair [14]. Notably, emerging evidence suggests that fibrocytes exhibit mesenchymal stem cell-like properties, raising the possibility that they contribute not only to matrix deposition but also to broader regenerative processes, including multipotent differentiation and tissue remodeling [15].

While the role of fibrocytes in tissue repair and fibrosis has been widely studied, their potential as a primary cell source for reprogramming into induced pluripotent stem cells (iPSCs) remains largely unexplored. Furthermore, their collection and use for this purpose in species such as cattle have not been previously attempted. Unlike traditional sources such as skin fibroblasts or embryonic cells, fibrocytes offer a distinct advantage as a readily accessible, diploid somatic cell population that can be isolated from peripheral blood. This enables minimally invasive collection, making fibrocytes an ideal cell source for field applications where aseptic sampling is critical. Compared to skin biopsies, which require specialized handling and pose a higher risk of microbial contamination, blood can be collected via simple venipuncture and either immediately processed or transported for culture under controlled conditions, allowing for the establishment of a stable cell population suitable for reprogramming.

Building on our previous work in generating and sustaining bovine iPSCs from fibroblasts [16,17], we aim to explore the feasibility of using fibrocytes as an alternative source for iPSC reprogramming. The ability to derive iPSCs from fibrocytes holds particular significance for livestock and wildlife species, as it enables the collection of live somatic cells from adult animals, facilitating broader applications in genetic preservation, reproduction and biotechnology. In this study, we establish protocols for the isolation and propagation of peripheral blood-derived fibrocytes from Holstein cattle, conduct transcriptomic characterization to evaluate their functional gene expression profiles in comparison to fibroblasts, and evaluate their potential for iPSC reprogramming.

## Materials and methods

### Animals, blood collection and preparation

Whole blood was collected by venipuncture of the caudal vein of Holstein cattle (*Bos taurus*; 1-2 years of age) in heparinized Vacutainer® tubes (BD, Switzerland) at the Cornell University Ruminant Center. Blood was processed immediately within 1-2 hours of collection for red blood cell lysis to obtain nucleated cells. For this step, blood was diluted in ammonium-chloride-potassium (ACK) red blood cell lysis buffer [composition: 150mM NH_4_Cl, 10mM KHCO_3_ and 0.1mM Na_2_EDTA] at a 1:9 ratio and incubated at 37°C for 5 minutes. The sample was then centrifuged for 5 minutes at 200 *x g* and supernatant removed. This process was repeated once to ensure the resultant pellet was devoid of red blood cells and appeared completely white. The pellet was then resuspended in phosphate buffered saline (PBS) and centrifuged again as a wash. This final pellet was plated in 0.1% gelatin coated dishes for cell culture (Figure 1A). Animal procedures were approved by the Institutional Animal Care and Use Committee of Cornell University. All chemicals were purchased from Millipore-Sigma (St. Louis, MO), unless otherwise noted.

**Figure 1.**
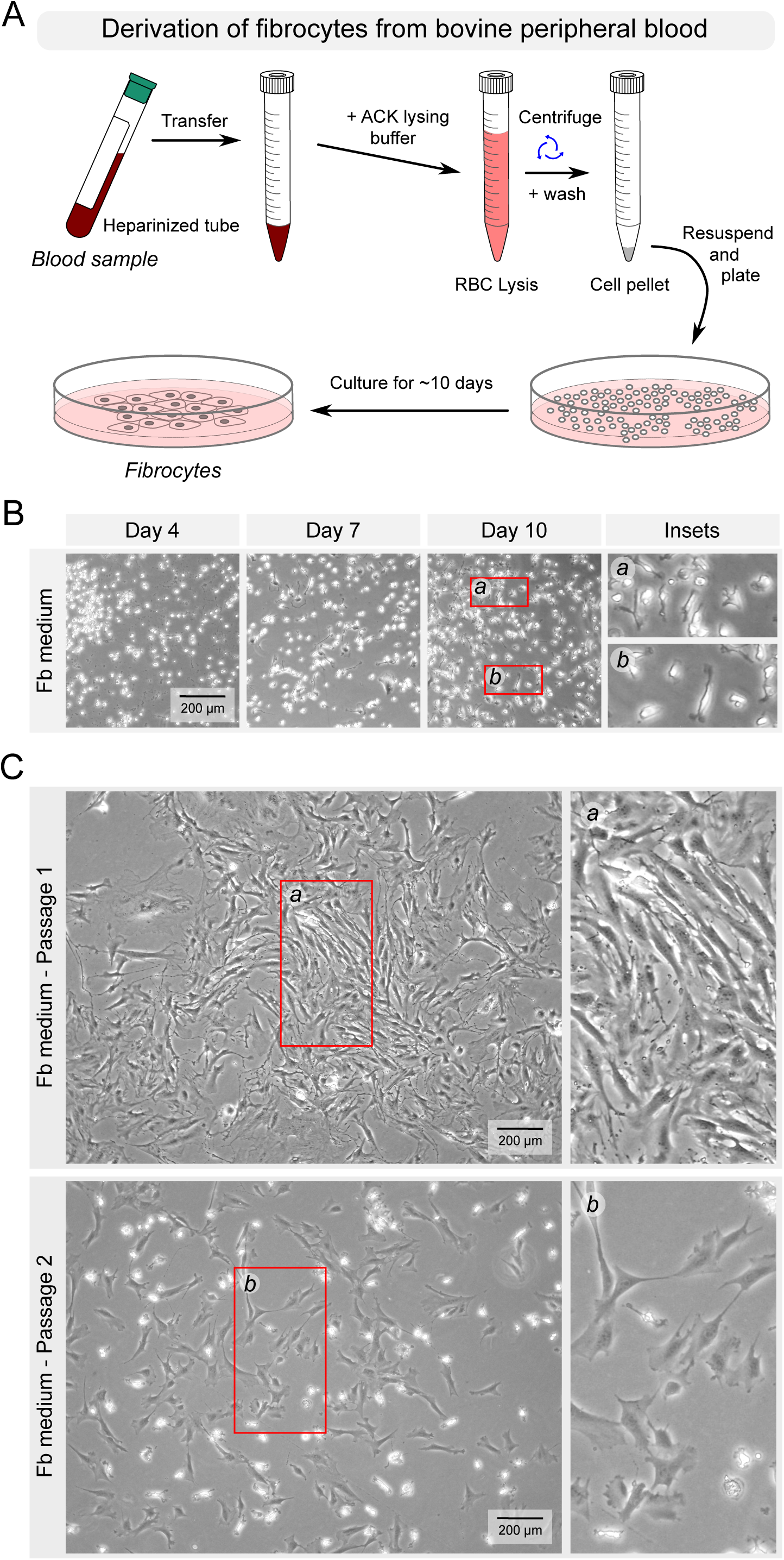
Derivation of fibrocytes from bovine peripheral blood and morphological characterization. (A) Workflow outlining the derivation of fibrocytes from bovine peripheral blood. Whole blood collected from the caudal vein was subjected to red blood cell lysis and washing steps. The resulting nucleated cell fraction was plated on gelatin-coated dishes in fibroblast (Fb) medium. Non-adherent cells were removed after 48 h, and adherent cells were expanded with daily medium replacement. Cultures were maintained until fibrocyte growth became evident and continued for approximately 10 days before the first passage. (B) Representative phase-contrast images showing progressive cellular changes during early culture in Fb medium. At Day 4, the culture consisted predominantly of small, phase-bright round cells with limited adherence. By Day 7, adherent micro-colonies emerged, and elongated spindle-shaped cells began to appear among residual round cells. By Day 10, two distinct morphologies were observed: rounded, refractile cells with smooth borders, and elongated bipolar cells with tapered ends and thin cytoplasmic extensions. (C) Morphological progression during passaging in Fb medium. Passage 1 cultures exhibited dense growth of elongated spindle-shaped cells aligned in parallel arrays and whorls. Passage 2 cultures at a lower density displayed greater heterogeneity, with stellate and triangular cells possessing broad lamellipodia and fine filopodial extensions interspersed among slender bipolar cells.

### Fibrocyte cell culture

The pellet of nucleated blood cells was resuspended in either a simple medium routinely used for culturing bovine fibroblasts [16]: Fibroblast medium [Fb; composition: Dulbecco’s High Glucose DMEM, 20% fetal bovine serum, penicillin-streptomycin, and non-essential amino acids (Gibco)], or in a modified mesenchymal stem cell medium: Fibrocyte medium [FbC; composition: Dulbecco’s High Glucose DMEM, 20% fetal bovine serum, penicillin-streptomycin, non-essential amino acids, 10nM dexamethasone, 0.1mM L-ascorbic acid-2-phosphate, 10ng/mL PDGF-bb(GoldBio), 10ng/ml EGF (GoldBio), 0.5µM A83-01(Cayman Chemical Company), and 1.5µM CHIR99021(Cayman Chemical Company)]. Resuspended cells were plated on 0.1% gelatin-coated cell culture dishes and incubated at 37°C in an atmosphere of 5% CO_2_. After 48 hours, the plate was gently washed with PBS to remove non-adherent cells and debris, and fresh medium was provided. Medium was subsequently replaced every 24 hours thereafter and observed for colony formation. Once plates were ∼70% confluent, colonies were passaged for experimentation or frozen for future use. Freezing was performed by resuspending cell pellets in 1 ml freezing medium (90% FBS with 10% DMSO), transferring to cryovials and placing them in –80°C in freezing containers (Thermo Scientific). After 24 hours, vials were moved to liquid nitrogen tanks for long-term storage. Matched samples from the same collection from each cow were cultured in Fb medium and FbC medium for comparing the different characterizations.

### Proliferation and morphological analysis

Fibrocyte morphology was assessed throughout culture using phase-contrast microscopy. Cells were seeded on 0.1% gelatin-coated culture dishes and maintained under standard conditions. Images were acquired at defined time points using a DM IL LED inverted microscope equipped with an MC190 HD camera (Leica Microsystems). Cell shape, spreading, and adherence characteristics were recorded, and representative fields were imaged at multiple magnifications to document progressive changes in morphology. Brightfield imaging was performed using the Lumaview 720/600 series live-imaging microscope to capture videos of cells cultured from initial peripheral blood draws following red blood cell lysis treatment. The recordings were used to visualize nucleation sites and monitor the evolving cellular morphology and proliferation dynamics over time, with individual image frames subsequently stitched together to produce a continuous time-lapse video. Cell proliferation assays documenting percent cell coverage were quantified from calibrated images acquired using a ZenCELL Owl live-cell imager (InnoME GmbH). Images were acquired from randomly selected fields within 24-well plates, in which four wells were allocated to each media condition (Fb or FbC) for each of two individual cows. Bright-field images were acquired every 10 minutes in a 24-well plate using a 10X magnification setting on the live-cell imager. The onboard algorithm segments each image by classifying pixels as either “cell-covered” or “background” based on contrast thresholds and then computes cell coverage (%) as the ratio of cell-covered pixels to total field area (overlapping cells are not double-counted).

For each cow, four wells per media condition (Fb and FbC) were imaged continuously. The cell coverage (%) values were plotted against time to generate growth curves for each well and media condition. For visualization and comparison, the median cell coverage across the four replicate wells per condition was plotted as the representative curve. The doubling time (T(d)) was determined by selecting the most representative exponential-growth segment of each curve and applying the standard proliferation-rate formula:

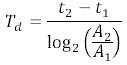

Where *A*_1_ and *A*_2_ represent cell coverage at times *t*_1_ and *t*_2_, respectively. Finally, the resulting image sequence from each treatment group was processed and compiled into continuous time-lapse videos to visualize cellular dynamics.

### RNA extraction and sequencing

Total RNA was extracted using the RNAqueous Micro Total RNA Isolation Kit (Thermo Fisher Scientific) using samples from three individual cows. For each animal, cells were cultured for 10 days at an equivalent seeding density under Fb or FbC medium conditions. Following this period, cells were passaged once, before RNA extraction. RNA quantity and purity were assessed by spectrophotometry (Nanophotometer N60, Implen), and integrity was confirmed using the Bioanalyzer 2100 (Agilent Technologies). Samples with RNA integrity numbers (RIN) > 8.0 were used for library preparation. RNA-Seq libraries were generated using the TruSeq Stranded mRNA Library Preparation Kit (Illumina), incorporating poly-A selection to enrich for mRNA. Libraries were quantified by qPCR and pooled for sequencing on an Illumina NovaSeq X Plus platform to obtain 150 bp paired-end reads. Read quality was evaluated with FastQC (Babraham Bioinformatics). Clean reads were aligned to the Bos taurus reference genome (ARS-UCD1.2) using STAR, and gene-level counts were obtained with EdgeR.

### Transcriptome analysis

DESeq2 was used for count normalization, and differential expression was performed using pair-wise quasi-likelihood comparisons in edgeR, and differentially expressed genes (DEGs) were identified using pairwise comparisons with an FDR < 0.05. For comparative analyses involving primary fibroblasts, publicly available RNA-seq data from NCBI GEO (GSE61027) [17] were incorporated and analyzed in parallel using the same normalization and statistical thresholds. Canonical pathway and upstream regulator analyses were performed using Ingenuity Pathway Analysis (IPA, Qiagen). For fibrocyte-fibroblast comparisons, canonical pathways with activation z-scores ≥ +2 or ≤ −2 were classified as activated or inhibited and grouped into functional biological categories. Pathways were visualized using bubble plots, to identify those with strongly activated or suppressed z-scores. Upstream regulator analysis identified transcription factors and receptors predicted to drive fibrocyte-specific gene expression programs; regulators with |z| ≥ 2 and p < 0.05 were retained. Normalized counts per million (CPM) for genes encoding these regulators were row-scaled and visualized using the pheatmap package with hierarchical clustering to illustrate predicted activation and inhibition patterns. To evaluate transcriptional responses attributable to the components of the FbC medium, DEGs contributing to the identification of dexamethasone, TGFB1, EGF, and WNT as upstream regulators were extracted and grouped according to regulator assignment. Shared and unique gene sets were determined by intersecting the DEG lists across regulators, yielding discrete categories representing convergent or factor-specific responses. Circos plots were generated using the circlize package to visualize the extent of transcriptional overlap among these regulator-defined gene sets.

### Reprogramming to pluripotency

Reprogramming of bovine fibrocytes was performed using methods previously described [18], with minor modifications. Briefly, fibrocytes were plated (5 x 10^4^ cells per well) and infected with a human polycistronic lentiviral vector encoding OCT4, SOX2, KLF4, and c-MYC [19] [18], together with a lentiviral SV40 Large T antigen vector [18]. Lentiviral packaging and production were carried out as described previously [16]. Medium replacement and transfer to irradiated mouse embryonic fibroblast (iMEF) feeders were carried out as indicated. Cultures were then continued in GMTi medium [DMEM/F12 containing N2 supplement, B–27 supplement, 1% non-essential amino acids supplement, 1% penicillin-streptomycin, 0.1 mM β-mercaptoethanol, 1.5 μM CHIR99021, 1 μM PD0325901, 0.5 μM A83–01, and 20 ng/ml hLIF] for colony expansion and maintenance [18]. Colonies with compact morphology, high nuclear-to-cytoplasmic ratios, and defined borders were manually picked and expanded for characterization.

### Alkaline phosphatase detection

Labeling for alkaline phosphatase (ALP) activity was performed as previously described [16]. Briefly, ALP activity was visualized using the Vector Blue AP Substrate Kit (Vector Laboratories) according to the manufacturer’s instructions. Reagents were mixed in 10 mL Tris–HCl buffer (pH 8.5) and added directly to cell culture plates, followed by incubation for 30 min at 37 °C in 5% CO_2_. Stained colonies were imaged under transmitted light without contrast using a DM IL LED inverted microscope equipped with an MC190 HD camera (Leica Microsystems).

### Reverse transcription polymerase chain reaction

Complementary DNA (cDNA) was synthesized from 2 µg of total RNA using the High-Capacity cDNA Reverse Transcription Kit with MultiScribe™ Reverse Transcriptase (Life Technologies) and Oligo(dT) primers (Bio Matik). Reverse transcription was performed in a C1000 Touch thermal cycler (Bio-Rad) under the following conditions: 25°C for 10 min (primer annealing), 37°C for 120 min (cDNA synthesis), and 85°C for 5 min (enzyme inactivation), followed by a hold at 4°C. The resulting cDNA was diluted 1:2 with nuclease-free water and stored at −20 °C until further use. Gene-specific primers (Table S1) were subsequently used for PCR amplification of bovine targets. Each 20 µL PCR reaction contained 1 µL of cDNA template, 0.4 µL of 10 mM dNTPs, 1 µL each of 10 µM forward and reverse primers, 0.5 µL of Taq polymerase, 2 µL of 10× PCR buffer, and nuclease-free water to volume.

## Results

### Derivation of fibrocytes from bovine peripheral blood

To establish a reproducible method for deriving fibrocytes from adult cattle, nucleated cells obtained from 10 ml peripheral blood were cultured under adherent conditions in fibroblast medium (Fb) (Figure 1). After rinsing off non-adherent cells at 48 hours, the remaining population was maintained under daily Fb medium replacement. During the first four days, cultures were dominated by small, phase-bright round cells showing minimal adherence. By Day 7, an increasing number of adherent cells formed discrete micro-colonies, and elongated spindle-shaped cells began to appear among residual round cells (Figure 1B). By Day 10, two prominent morphotypes were consistently observed: rounded, refractile cells with smooth contours and elongated bipolar cells with tapered ends and fine cytoplasmic extensions. These morphological transitions were indicative of a progressive adaptation of adherent leukocytes toward a fibroblast-like phenotype consistent with fibrocyte differentiation. Time-lapse imaging captured this process in real time (Movie S1), showing fibrocyte emergence from a colony-forming nucleation site, followed by outward proliferation and migration across the gelatin-coated surface.

Upon passaging, cultures demonstrated further morphological stabilization. Passage 1 cells proliferated rapidly, forming dense formations of elongated spindle-shaped cells organized in parallel streams and whorled arrangements (Figure 1C). At lower seeding density (Passage 2), cultures became more heterogeneous, displaying stellate and triangular cells with broad lamellipodia and filopodial processes interspersed among slender bipolar cells. Together, these observations confirmed successful derivation and propagation of fibrocytes from bovine peripheral blood under standard Fb culture conditions.

### Distinct functional identity of bovine fibrocytes

To define the molecular characteristics that distinguish fibrocytes from fibroblasts, we performed RNA-seq analysis followed by principal component analysis (PCA) and differential expression profiling. PCA revealed clear segregation of the two cell types along the first principal component, which captured the majority of gene expression variance (Figure 2A). Samples from three independent animals clustered tightly within each group, indicating high within-group consistency and robust separation between fibrocyte and fibroblast transcriptomes.

**Figure 2.**
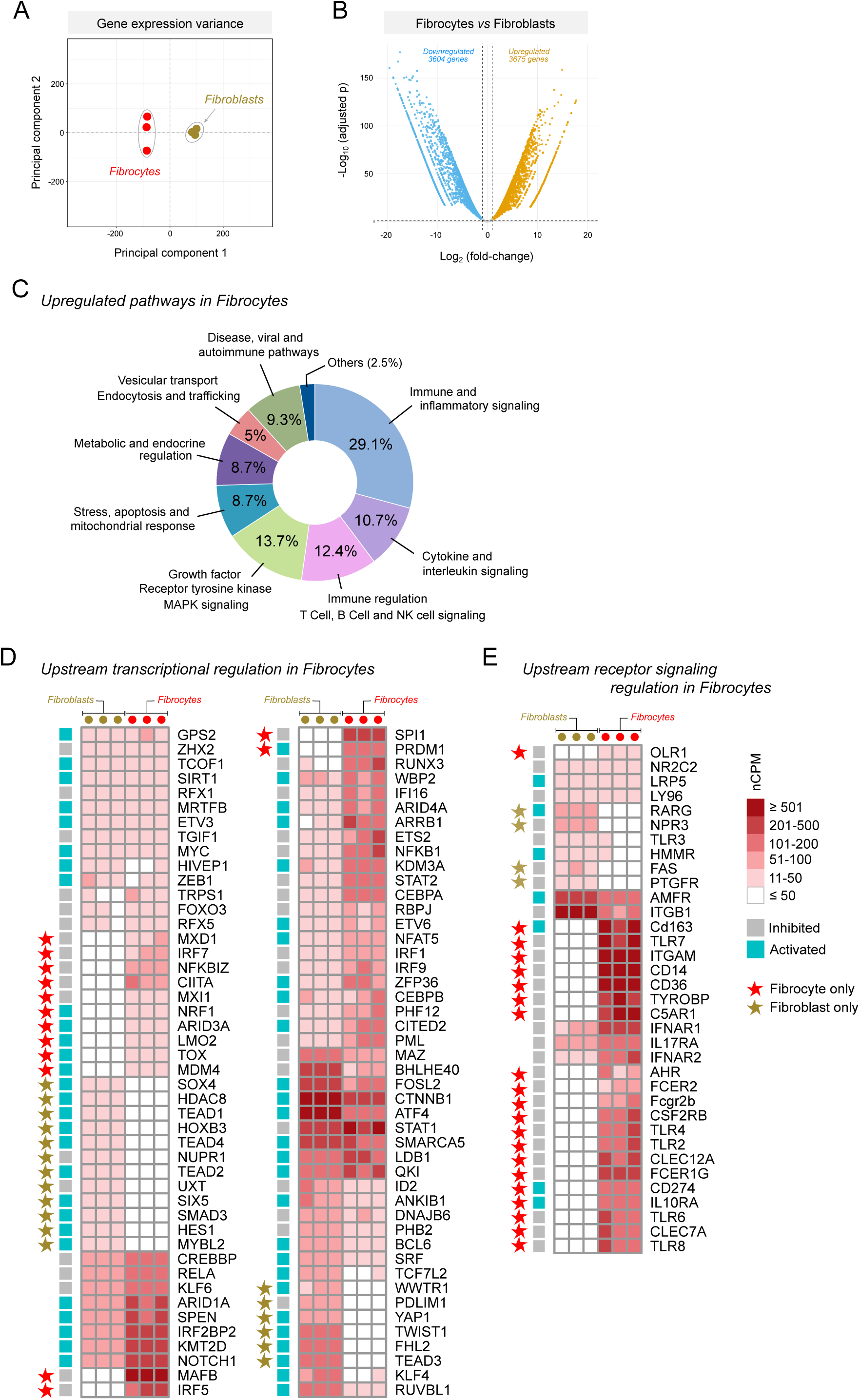
Transcriptomic distinction and functional enrichment of fibrocytes compared to fibroblasts. (A) Principal component analysis (PCA) of RNA-seq data showing separation of fibrocytes and fibroblasts based on global gene expression variance. Each point represents an independent animal sample. (B) Volcano plot comparing differential gene expression between fibrocytes and fibroblasts. A total of 3,675 genes were significantly upregulated and 3,604 genes were downregulated in fibrocytes (adjusted p < 0.05). (C) Functional enrichment analysis of genes upregulated in fibrocytes identifying major biological processes and pathways. Prominent categories included immune and inflammatory signaling, cytokine and interleukin signaling, growth factor–mediated MAPK signaling, metabolic and endocrine regulation, and stress and mitochondrial response pathways. (D) Upstream transcriptional regulators differentially expressed between fibrocytes and fibroblasts. Heatmaps represent normalized counts per million (nCPM) for key transcription factors predicted to drive fibrocyte-specific gene expression. (E) Upstream receptor signaling regulators in fibrocytes. Heatmap displays nCPM values for receptors and signaling molecules differentially expressed between fibrocytes and fibroblasts, highlighting pathways related to immune recognition, cytokine and growth factor signaling, and cellular communication. For both (D) and (E), regulators uniquely expressed in fibrocytes or fibroblasts are indicated by different stars. Color-coded boxes indicate predicted activation or inhibition status of the upstream regulation.

Differential expression analysis identified extensive remodeling of the fibrocyte transcriptome, with 3,675 genes significantly upregulated and 3,604 genes downregulated relative to fibroblasts (adjusted p < 0.05; |log_2_ fold-change| ≥ 1; Figure 2B). Functional enrichment of the upregulated genes revealed dominant representation of immune and inflammatory signaling networks (29.1%), including cytokine, interleukin, and interferon pathways (10.7%), as well as T-, B-, and NK-cell signaling modules (12.4%) (Figure 2C). Additional enriched categories included growth factor–mediated MAPK signaling (13.7%), metabolic and endocrine regulation (8.7%), vesicular trafficking and endocytosis (5%), stress– and mitochondria-related response pathways (8.7%), and Disease, viral and autoimmune pathways (9.3%). These findings indicate that fibrocytes maintain a hybrid transcriptional program integrating immune-effector and mesenchymal regulatory functions.

Upstream regulator analysis revealed coordinated activation of transcriptional and receptor-mediated signaling networks that define fibrocyte identity (Figure 2D,E). Among transcriptional regulators, a fibrocyte-restricted set, comprising SPI1 (PU.1), IRF5, IRF7, NFKBIZ, PRDM1 (BLIMP1), CIITA, MXD1, MXI1, NRF1, MAFB, ARID3A, LMO2, TOX, and MDM4, was predicted to drive key fibrocyte-enriched gene modules. These regulators collectively represent inducible inflammatory and antiviral control (IRF5/IRF7/NFKBIZ/SPI1), antigen-presentation potential (CIITA), hematopoietic lineage memory (ARID3A/LMO2/TOX), and metabolic-stress adaptation (MXD1/MXI1/NRF1/MAFB).

Complementing this transcriptional framework, receptor-level analyses identified a coordinated upregulation of immune and cytokine receptors, including TLR2, TLR4, TLR6, TLR7, TLR8, FCER1G/2, IL10RA, CD14, CD163, C5AR1, and CD274, as well as adhesion and signaling molecules such as ITGAM, CD36, TYROBP, and FAS (Figure 2E). Together, these receptor and transcriptional networks define a fibrocyte-specific regulatory system that integrates immune sensing, stress adaptation, and reparative signaling, distinguishing fibrocytes as a molecularly distinct somatic lineage rather than a fibroblast variant. All DEGs and curated gene lists corresponding to each enrichment, upstream regulator, and receptor analysis shown in Figure 2 are provided in Supplementary Data File 1.

### Fibrocyte medium (FbC) promotes stable propagation of blood-derived fibrocytes

Because fibrocytes share features of both mesenchymal and hematopoietic lineages, we developed a modified culture formulation, the fibrocyte medium (FbC), designed to promote proliferation while maintaining cellular plasticity. FbC incorporates dexamethasone, ascorbic acid, PDGF-BB, EGF, and small-molecule inhibitors (A83-01 and CHIR99021) to support both mesenchymal expansion and inhibit immune activation that might trigger spontaneous senescence. When nucleated peripheral blood cells were cultured in FbC, small, phase-bright round cells initially dominated the culture. By Day 7, adherent micro-colonies emerged, and by Day 10, elongated spindle-shaped fibrocytes became prevalent, displaying organized parallel alignment (Figure 3A). Upon passaging, fibrocytes retained a uniform spindle morphology and proliferated rapidly with consistent growth across passages, indicating that FbC medium supported a stable and self-sustaining fibrocyte population (Figure 3B).

**Figure 3.**
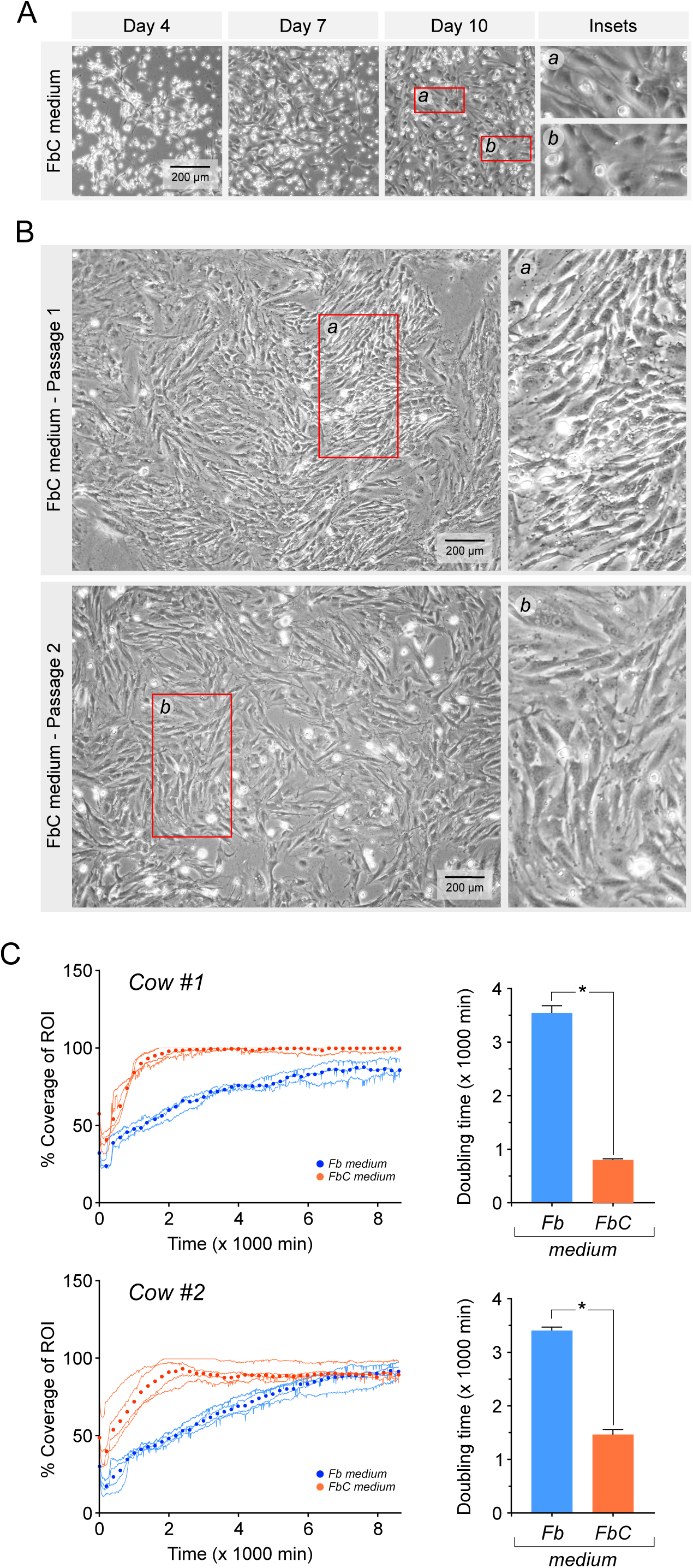
Derivation and growth kinetics of bovine fibrocytes cultured in fibrocyte medium (FbC). (A) Sequential phase-contrast images showing the emergence of fibrocytes from bovine peripheral blood in FbC medium. Small, phase-bright round cells initially dominated the culture, giving rise to adherent micro-colonies by Day 7 and elongated spindle-shaped fibrocytes by Day 10, indicative of progressive differentiation and adaptation to adherent growth. (B) Representative morphology of passaged fibrocytes maintained in FbC medium showing uniform spindle shape, organized parallel alignment, and sustained proliferation across passages, consistent with stable expansion under these conditions. (C) Growth kinetics comparing fibrocytes cultured in Fb and FbC media. Cultures in FbC exhibited a significantly higher proliferation rate and shorter population-doubling time, indicating that FbC medium supports more robust fibrocyte expansion. Traces in the growth curve represent independent replicates, and data points indicate median values. Bar graphs show mean ± SEM from three independent biological replicates (*p < 0.05).

To visualize these differences dynamically, time-lapse imaging was used to capture fibrocyte behavior under both culture conditions. In Fb medium, fibrocytes exhibited diffuse growth, extensive migration, and morphological heterogeneity characterized by broad lamellipodia and frequent shape transitions (Movie S2). In contrast, fibrocytes cultured in FbC medium displayed reduced motility, earlier attachment, and coordinated elongation, followed by rapid mitotic activity and organized palisading proliferation (Movie S3). These live-cell recordings underscore the stabilizing and proliferative effects of FbC medium, revealing a shift from migratory adaptation in Fb to sustained, patterned growth in FbC.

Quantitative growth kinetics further substantiated these observations in that fibrocytes cultured in FbC medium proliferated significantly faster than those maintained in Fb medium (Figure 3C). FbC medium cultures exhibited steeper exponential growth trajectories and significantly shorter population-doubling times (p < 0.05), reflecting enhanced proliferative capacity. These findings demonstrate that the FbC formulation markedly improves fibrocyte proliferation while maintaining a mesenchymal-like, plastic phenotype suited for downstream applications.

### FbC medium factors cooperatively reprogram fibrocyte transcriptomes

To determine how FbC medium alters fibrocyte gene expression and functional state, we performed transcriptomic profiling of fibrocytes cultured in Fb versus FbC medium conditions. Principal component analysis revealed separation of samples by medium type, indicating a reproducible transcriptional shift induced by FbC formulation (Figure 4A). Differential expression analysis identified 2,081 genes that were significantly altered between the two conditions (adjusted p < 0.05, |log_2_FC| > 1), with 1,140 genes upregulated and 941 genes downregulated in fibrocytes cultured in FbC medium (Figure 4B). Hierarchical clustering further confirmed distinct expression signatures, with consistent segregation of FbC-treated fibrocytes across biological replicates (Figure 4C).

**Figure 4.**
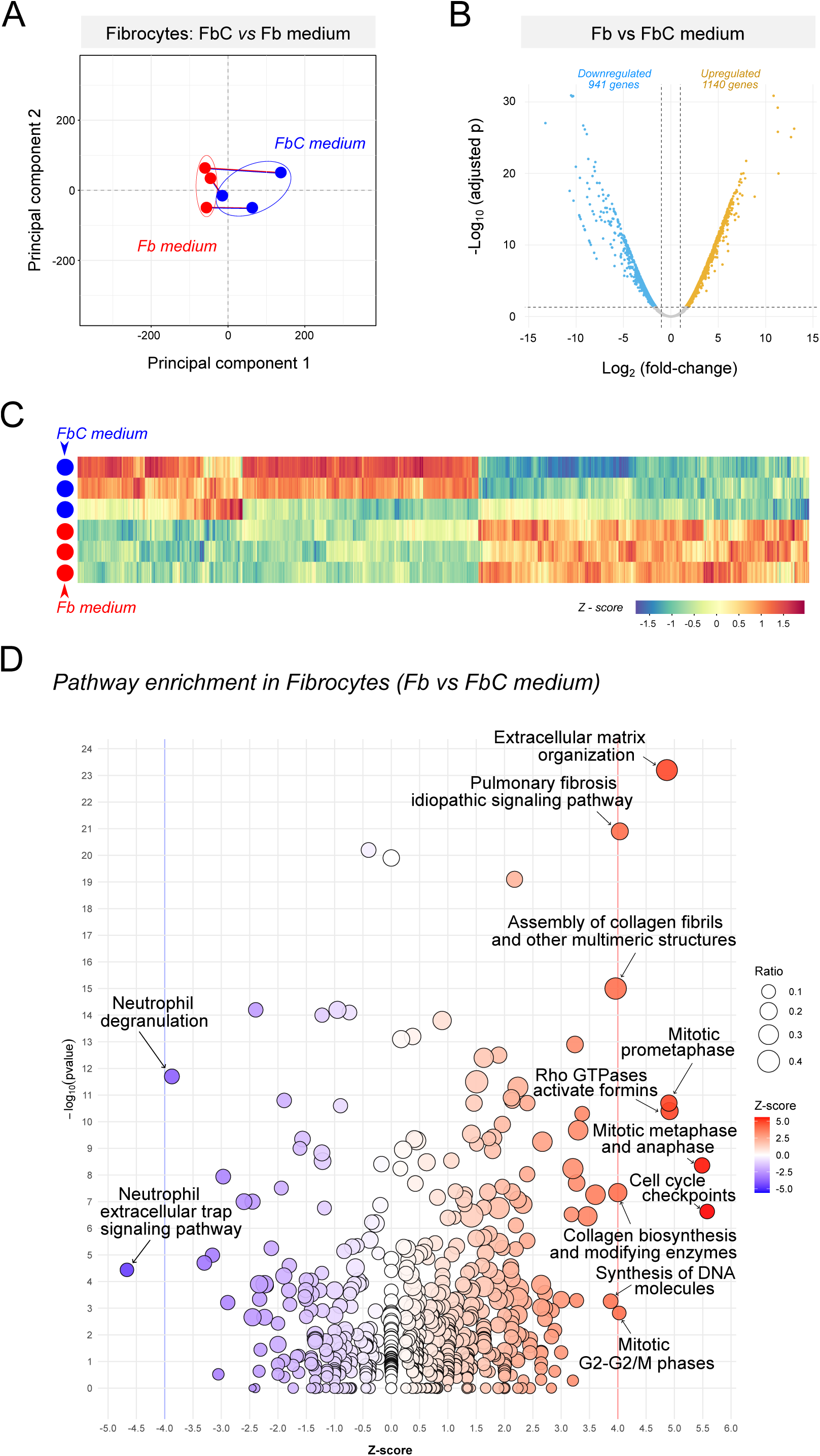
Transcriptomic effects of FbC medium on fibrocyte gene expression. (A) Principal component analysis (PCA) showing distinct clustering of fibrocyte transcriptomes cultured in FbC medium (blue) versus Fb medium (red), indicating a strong global transcriptional shift driven by medium composition. (B) Volcano plot of differential gene expression between Fb and FbC conditions identifying 1,140 upregulated and 941 downregulated genes in fibrocytes cultured in FbC medium (adjusted p < 0.05, |log FC| > 1). (C) Heatmap of differentially expressed genes (Z-score normalized) illustrating consistent transcriptional remodeling of fibrocytes across biological replicates in response to FbC medium. (D) Pathway enrichment analysis of differentially expressed genes showing activation of proliferative, matrix assembly, and cell-cycle pathways (e.g., mitotic progression, collagen biosynthesis, Rho GTPase signaling) in FbC medium, with concurrent downregulation of inflammatory response programs. Together, these data demonstrate that FbC medium promotes a proliferative, matrix-constructive, and reparative transcriptional state in fibrocytes relative to Fb medium.

Pathway enrichment analysis highlighted a broad reorganization of cellular programs associated with proliferation, cytoskeletal remodeling, and extracellular matrix assembly (Figure 4D). Upregulated pathways included collagen biosynthesis and modification, assembly of collagen fibrils, mitotic cell-cycle progression, DNA replication, and Rho GTPase-regulated cytoskeletal dynamics, indicative of enhanced matrix production and proliferative activity. Conversely, pathways associated with inflammatory signaling were significantly downregulated, suggesting suppression of immune effector functions.

Upstream regulator-driven gene signatures revealed that each FbC component, dexamethasone, TGFB1, EGF, and WNT, exerted a distinct yet highly interconnected transcriptional influence on fibrocytes (Figure 5A). Dexamethasone showed the broadest influence (444 genes), followed by TGFB1 (424 genes), EGF (160 genes), and WNT activation (32 genes). Dexamethasone, TGFB1, and EGF formed the dominant axis of overlap, but WNT displayed dense connectivity to the other three pathways; only 203, 176, 24, and 5 genes, respectively, were unique to each regulator. Canonical pathway analysis on the genes responsive to the four signaling inputs revealed strong activation of proliferative and matrix-constructive pathways (Figure 5B). Upregulated pathways included mitotic cell-cycle checkpoints, extracellular matrix organization, collagen biosynthesis and fibril assembly, Rho GTPase–formin signaling, and multiple growth factor–responsive modules such as MET, PDGF, and GP6 signaling. In contrast, several immune and inflammatory pathways were consistently predicted to be suppressed. These included IL-17F signaling, NOD1/2 signaling, macrophage classical activation, neutrophil degranulation, TREM1 signaling, and neutrophil extracellular trap formation. These changes indicated that the FbC medium attenuates inflammatory activation while favoring reparative and proliferative fibrocyte states. All DEG tables and curated gene sets corresponding to the medium comparison and the individual FbC component analyses shown in Figures 4 and 5 are provided in Supplementary Data File 2.

**Figure 5.**
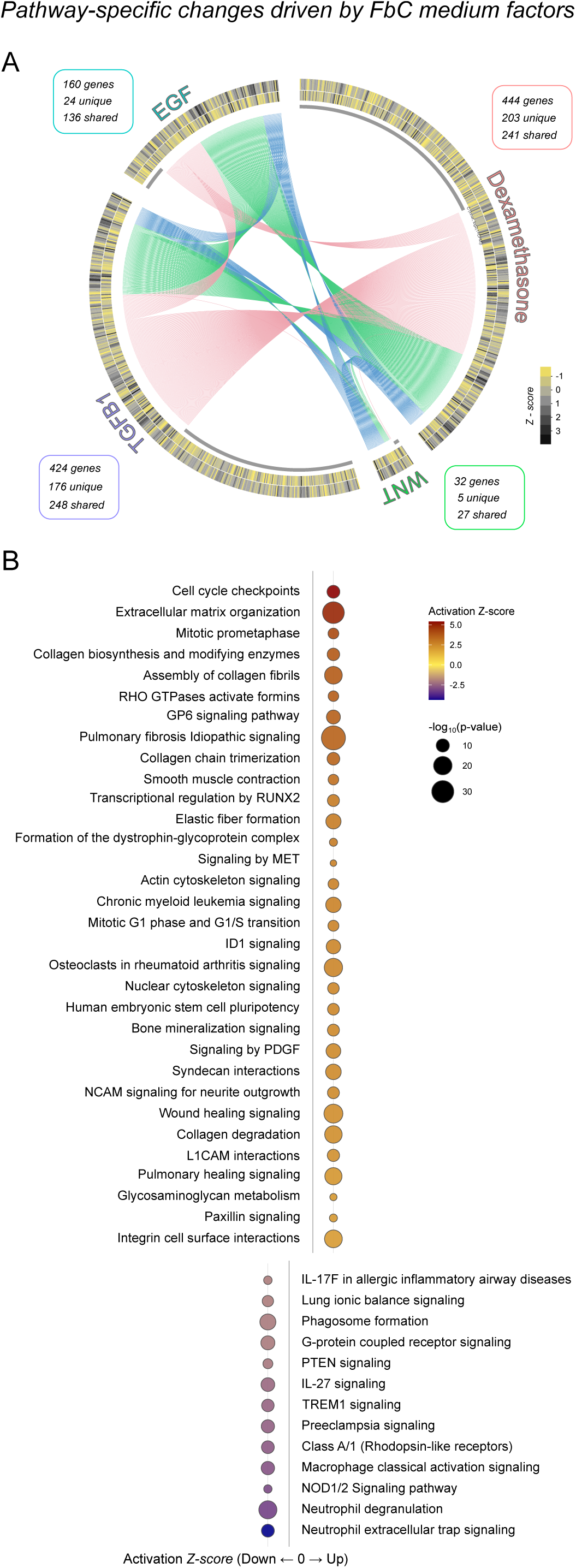
Pathway-specific transcriptomic modulation by FbC medium factors. Circos diagram depicting the extent and overlap of gene expression changes induced by dexamethasone, TGFB1, EGF, and WNT activation in bovine fibrocytes cultured in FbC vs Fb. Outer tracks show scaled expression values (Z-scores) for all differentially expressed genes associated with each factor. Chord connections indicate shared gene responses, with color coding proportional to the degree of overlap: pink for genes shared by two additives, green for genes shared by three, and blue for genes common to all four additives. Dexamethasone regulated the largest gene set, followed by TGFB1, EGF and WNT. (B) Canonical pathway enrichment analysis showing pathways significantly affected by dexamethasone, TGFB1, EGF, and WNT signaling in fibrocytes. Bubble size corresponds to –log_10_(p-value), and bubble color denotes activation state. The combined actions of these factors strongly activated proliferative and matrix-constructive programs, while suppressing inflammatory and innate-immune pathways.

### Fibrocyte-specific functional gene networks reveal integrated immune and reparative programs

To define the molecular architecture underlying fibrocyte identity, we mapped genes uniquely expressed in fibrocytes cultured in both Fb and FbC media but largely absent or undetectable in fibroblasts. High-confidence protein–protein interaction networks were constructed and curated into functional themes reflecting coherent biological processes (Figure 6). Distinct, interconnected modules emerged, revealing that fibrocytes integrate immune signaling, antigen-processing, and reparative programs within a unified transcriptional framework. Prominent networks included chemokine and cytokine receptors, NF-κB/MAPK modulators, and GPCR/lipid mediator signaling, reflecting the capacity for dynamic inflammatory and paracrine communication. Additional clusters representing antigen presentation and processing, complement system, and costimulatory or immune-synapse components highlighted fibrocytes’ immune-regulatory potential and antigen-presenting capacity. Modules associated with monocyte/scavenger identity and efferocytosis, together with adhesion/integrin signaling and pattern-recognition receptors, underscored fibrocytes’ phagocytic and matrix-interactive functions. A prominent transcriptional regulation cluster encompassing multiple immune regulators linked to Type I interferon and antiviral response networks suggested a strong innate defense axis characteristic of leukocyte/monocyte-derived populations. Finally, smaller but consistent modules related to DNA repair and genome stability implied sustained proliferative potential and resilience under culture adaptation.

**Figure 6.**
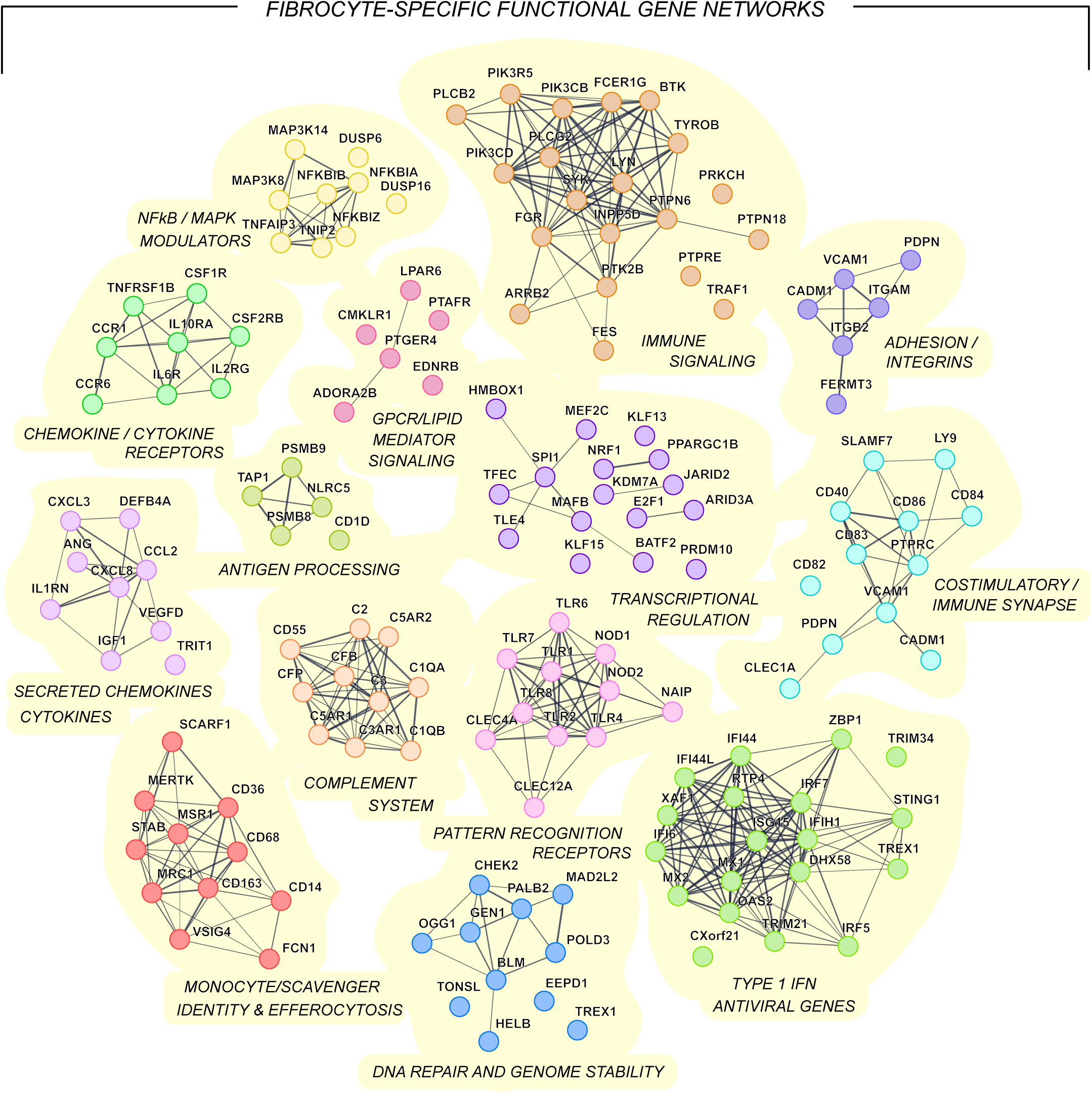
Fibrocyte-specific functional gene networks. Functional gene networks delineated from genes uniquely expressed in fibrocytes cultured in both Fb and FbC media and largely absent in fibroblasts. High-confidence protein–protein interaction maps were generated and organized into curated functional themes to reveal the dominant molecular programs defining fibrocyte identity. Major network themes include chemokine and cytokine receptors, secreted cytokines, NF-κB/MAPK modulators, GPCR/lipid mediator signaling, antigen presentation and processing, complement system, monocyte/scavenger identity and efferocytosis, pattern-recognition receptors, Type I interferon and antiviral genes, transcriptional regulation, immune signaling, adhesion/integrins, costimulatory and immune-synapse components, and DNA repair/genome stability pathways. Together, these networks outline the coordinated immune-regulatory and reparative signature that distinguishes fibrocytes from fibroblasts.

### Induction of pluripotency in bovine fibrocytes

To determine whether bovine fibrocytes can be reprogrammed to a pluripotent state, fibrocyte cultures established in FbC medium were subjected to lentiviral transduction using a polycistronic OSKM+LT (Figure 7A). Following infection, fibrocytes were maintained under feeder-free conditions for one week before transfer onto irradiated MEF feeders and transition to GMTi medium to facilitate complete reprogramming. Distinct morphological transitions were observed during the reprogramming process (Figure 7B). By Day 7, fibrocytes formed somewhat dense, proliferative clusters of small, rounded cells that progressively condensed into tightly packed colonies by Day 15. These colonies displayed smooth, dome-shaped borders and high refractility, consistent with a pluripotent morphology. Upon passaging, established fibrocyte-derived colonies exhibited stable growth as compact, multilayered structures with defined colony edges and a high nucleus-to-cytoplasm ratio (Figure 7C). Alkaline phosphatase (ALP) staining confirmed robust activity in multiple colonies derived from independent cultures, indicative of successful reprogramming and acquisition of a pluripotent phenotype (Figure 7D).

**Figure 7.**
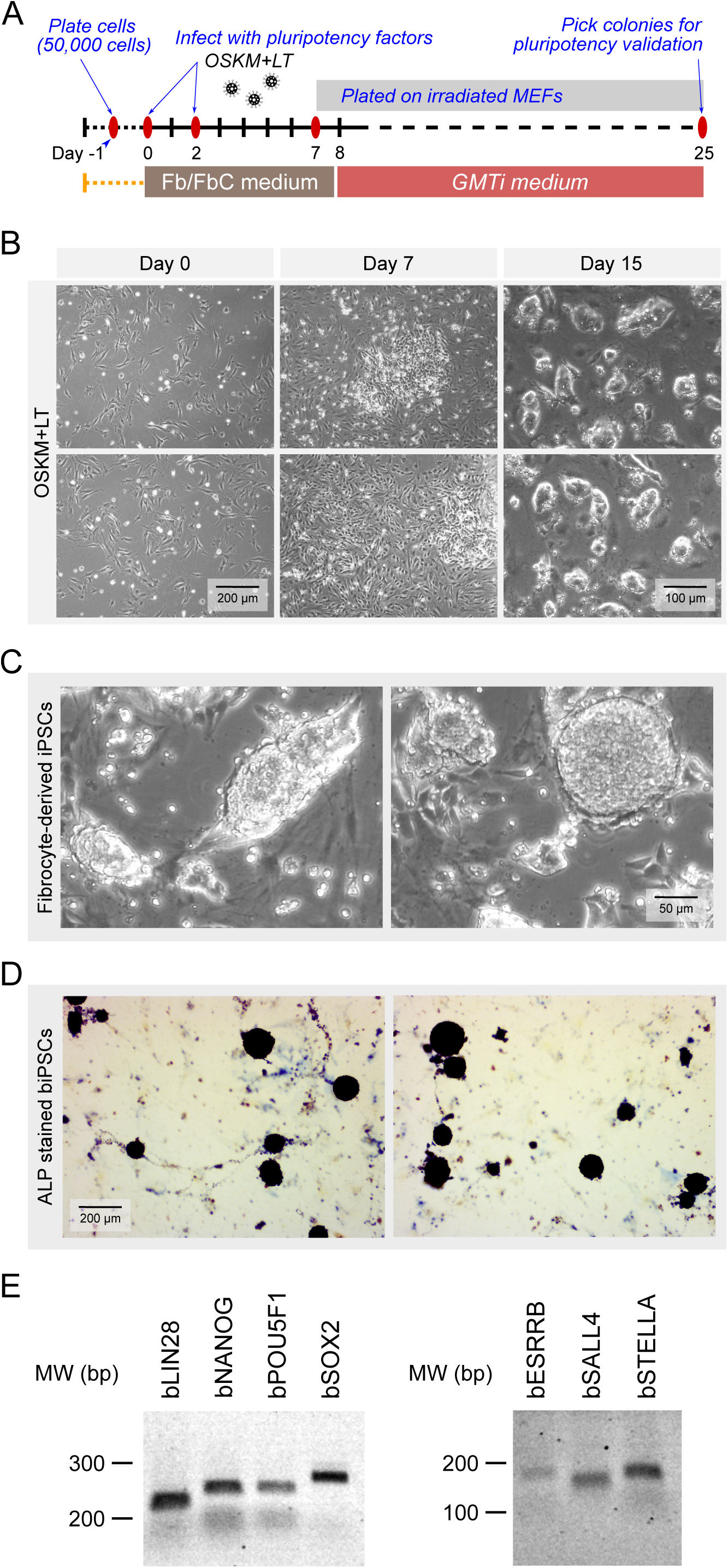
Induction of pluripotency in bovine fibrocytes. (A) Experimental timeline for reprogramming fibrocytes to induced pluripotent stem cells (iPSCs). Fibrocytes were plated (Day –1) in FbC medium, transduced twice with OSKM+LT (Days 0 and 2), transferred to irradiated MEF feeders (Day 7-8), switched to GMTi medium to facilitate reprogramming and sustenance. Putative colonies were picked for propagation and validation within Days 18-25. (B) Representative phase-contrast images showing progressive morphological changes during reprogramming of fibrocytes at Day 0, 7, and 15 after infection. Compact, dome-shaped colonies resembling iPSCs emerged by Day 15. Two representative replicate cultures are shown. (C) High-magnification views showing morphology of established fibrocyte-derived iPSC colonies after passaging with characteristic compact, refractile colony borders typical of bovine iPSCs. (D) ALP staining of biPSC colonies generated from fibrocytes, showing robust ALP activity in multiple colonies across two independent plates. Colonies displayed intense staining characteristic of pluripotent stem cells. (E) Endogenous pluripotency gene expression in fibrocyte-derived biPSCs by RT-PCR. Amplicons shown for core and naïve-associated markers (e.g., bovine POU5F1/OCT4, SOX2, KLF4, ESRRB, SALL4, and STELLA/DPPA3) with size ladders indicated (bp).

To validate endogenous activation of the pluripotency network, RT-PCR analysis was performed on fibrocyte-derived biPSCs (Figure 7E). Expression of core pluripotency markers POU5F1/OCT4, SOX2, and KLF4, as well as naïve-associated factors ESRRB, SALL4, and STELLA/DPPA3, were detected in all tested colonies. These results demonstrate that bovine fibrocytes, when cultured in optimized fibrocyte medium and reprogrammed under GMTi conditions, can give rise to bona fide iPSCs exhibiting both morphological and molecular hallmarks of pluripotency.

## Discussion

The recognition that circulating somatic cells can adopt reparative or mesenchymal-like phenotypes has fundamentally expanded our understanding of cellular plasticity in adult tissues [20,21]. Among these, fibrocytes have attracted attention as a rare leukocyte-derived population capable of migrating to sites of injury and contributing to extracellular matrix deposition, inflammation resolution, and tissue remodeling [22]. Despite their identification in multiple mammalian species, the precise molecular identity and lineage relationships of fibrocytes have remained ambiguous, largely due to the limitations of earlier descriptive studies and the absence of molecular profiling in non-rodent and non-human systems. In livestock, this gap has been particularly evident; no prior work had defined fibrocytes at the transcriptomic level or explored their potential as an accessible, field-adaptable somatic cell type. Understanding their molecular composition is therefore not only relevant to clarifying their biological role but also to assessing their suitability as a reprogrammable somatic cell source for pluripotency induction.

Fibrocytes exhibit a unique molecular and functional profile that bridges hematopoietic and mesenchymal lineages, reflecting their dual role in immune regulation and tissue repair [23]. Fibrocytes derived from bovine blood exhibit a similar molecular signature that reinforces this identity integrating immune, reparative, and metabolic programs. Upon inflammatory or fibrotic challenge, leukocyte-derived precursors traffic to sites of damage through chemokine-mediated recruitment involving CCR1– and CXCR4-associated signaling pathways [14,24–26], a migratory framework also reflected by chemokine and GPCR modules enriched in bovine fibrocytes. Within injured tissue, fibrocytes acquire a reparative phenotype characterized by type I collagen and fibronectin production [13,27], a program mirrored in bovine fibrocytes by the upregulation of COL1A1, COL3A1, FN1, MMP9, and SERPINE1 and the activation of collagen biosynthesis and fibril assembly pathways. Consistent with this reparative-immune duality, fibrocytes expressed high levels of IL6, TNF, and CCL2, reflecting conserved cytokine and chemokine responses that modulate local immune activity and coordinate monocyte and lymphocyte recruitment [28,29]. In parallel, fibrocytes exhibited coordinated upregulation of MHC II-associated and costimulatory genes (CD80, CD86), indicating that antigen-presenting potential, previously demonstrated in human fibrocytes [30], is conserved in the bovine lineage. Enrichment of growth-factor-mediated signaling networks, including activation of MAPK and PI3K pathways and increased expression of receptors such as CSF1R [31], further supports a role for paracrine responsiveness to cues such as TGF-β, PDGF, and VEGF in driving proliferative and angiogenic remodeling [32,33]. Moreover, prior evidence that fibrocytes can differentiate into osteogenic, chondrogenic, adipogenic and myogenic lineages [34,35], aligns with the metabolic and developmental plasticity observed in bovine fibrocytes. Collectively, these findings define fibrocyte derivation *in vitro* as an inducible extension of the hematopoietic lineage that integrates immune regulation, paracrine growth signaling, and tissue repair, features that provide a biological rationale for leveraging these cells as a plastic somatic platform for reprogramming.

Upstream regulator analysis identified a fibrocyte-restricted transcriptional network governed by SPI1, IRF5/IRF7, NFKBIZ, PRDM1, CIITA, and MAFB, defining a core regulatory axis characteristic of myeloid-derived cells. These transcription factors collectively delineate a hematopoietic control layer that persists in fibrocytes despite acquisition of mesenchymal and reparative features. SPI1 (PU.1) serves as a master regulator of myeloid specification, maintaining expression of receptors such as CSF1R, CD14, and integrins that underpin mononuclear phagocyte identity and immune competence [36,37]. The interferon regulatory factors IRF5 and IRF7 orchestrate inducible inflammatory and antiviral programs downstream of Toll-like receptor signaling [38,39], while NFKBIZ (IκBζ) acts as a noncanonical NF-κB co-activator that controls late-phase cytokine genes including IL6 and TNF [40,41]. Together, these regulators create an inducible immune–stress axis consistent with the cytokine and chemokine responses observed in fibrocytes. In contrast, PRDM1 (BLIMP1) and MAFB temper proinflammatory signaling and guide monocyte-to-macrophage maturation, reinforcing a transcriptional switch toward resolution and repair. PDRM1, through broadly expressed across hematopoietic lineages, functions in monocytes and macrophages as a transcriptional repressor that limits inflammatory cytokine production and promotes transition towards resolution [42]. MAFB, a transcription factor downstream of SPI1, promotes monocyte-to-macrophage differentiation and represses proliferative signaling while integrating lipid and anti-inflammatory responses [43]. Finally, CIITA, the master transactivator of MHC class II genes, provides a molecular basis for the antigen-presenting potential reflected in fibrocyte expression of MHC II, CD80, and CD86 [44]. The coordinated expression of these factors in bovine fibrocytes therefore supports a model in which partial hematopoietic lineage memory is transcriptionally retained.

Optimization of the fibrocyte culture environment was essential to sustain proliferation while preserving lineage plasticity. Conventional Fb medium supported only limited adherence and expansion, suggesting that the basal conditions failed to maintain the signaling equilibrium required for robust fibrocyte growth. In contrast, the FbC formulation – incorporating dexamethasone, ascorbic acid, PDGF-BB, EGF, A83-01, and CHIR99021, produced rapid adherence in fibrocyte derivation and markedly enhanced proliferation. Transcriptomic comparisons between Fb and FbC cultures revealed extensive factor-driven programming across cell-cycle, mitotic progression, and cytoskeletal remodeling, accompanied by suppression of inflammatory signaling. Glucocorticoid signaling is known to restrain proinflammatory activity through MAPK inhibition, NF-κB interference, and induction of the phosphatase DUSP1/MKP-1 [45], providing a mechanistic basis for the dampening of immune programs observed under FbC conditions. Ascorbate supports cellular remodeling and viability by promoting prolyl and lysyl hydroxylase activity and procollagen maturation, while also enhancing TET-dependent DNA demethylation that improves somatic cell reprogramming competence [46–49], consistent with the proliferative, plastic state we detect in FbC cultures. PDGF-BB activates PI3K–AKT and ERK pathways to drive mesenchymal stem cell proliferation, motility, and survival [50,51], aligning with the cell-cycle and cytoskeletal programs enriched in FbC relative to Fb. In parallel, soluble EGF augments human MSC proliferation while preserving early progenitors and boosting paracrine outputs [52], providing a contextual basis for the proliferative, pro-angiogenic signaling we observe in FbC cultures. To avoid TGF-β–driven differentiation, the Activin/NODAL/TGFβ pathway inhibitor A83-01 supports expansion of undifferentiated mesenchymal and epithelial progenitors and enhances reprogramming efficiency in somatic cells [53], consistent with the goal of maintaining a reprogrammable fibrocyte state. Finally, CHIR99021 stabilizes β-catenin to activate canonical WNT signaling, a well-established mechanism for sustaining self-renewal and cytoskeletal organization in mesenchymal stem cell systems [54,55], matching the FbC-induced transcriptional shift toward proliferative, motile phenotypes. Collectively, these component-level mechanisms explain the FbC-driven convergence on MAPK and PI3K activation with inflammatory restraint, yielding a stable growth-permissive equilibrium that preserves fibrocyte identity and plasticity.

The establishment of a stable, proliferative fibrocyte population under FbC conditions provided a biologically advantageous platform for somatic reprogramming. Fibrocytes possess intrinsic features that may lower the epigenetic barrier to pluripotency, including active chromatin states associated with inflammatory signaling, mitochondrial metabolism, and stress adaptation. These characteristics parallel those seen in hematopoietic or myeloid-derived cells that reprogram with relatively high efficiency compared to terminally differentiated fibroblasts [56,57]. The retention of hematopoietic transcriptional regulators such as SPI1, IRF5, and PRDM1, together with metabolic and proliferative signatures reinforced by FbC conditions, likely facilitates chromatin accessibility and responsiveness to the OSKM + LT factors used for reprogramming [18]. Consistent with this, fibrocytes cultured in FbC medium transitioned efficiently to compact, alkaline-phosphatase positive colonies that expressed endogenous POU5F1, SOX2, KLF4, and naïve pluripotency factors (ESRRB, SALL4, STELLA/DPPA3), demonstrating authentic induction of pluripotency. These observations align with prior evidence that optimized mesenchymal or hematopoietic contexts enhance iPSC derivation through PI3K–AKT and WNT signaling, as well as through vitamin C-driven epigenetic remodeling [58–60]. In this context, bovine fibrocytes represent a naturally accessible, field-adaptable somatic cell type that can be isolated aseptically from peripheral blood and stably propagated without genomic alteration. The ability to derive iPSCs from these cells extends the feasibility of creating genetically defined pluripotent lines from valuable or endangered livestock, establishing a foundation for genetic preservation, reproductive biotechnology, and regenerative applications in large-animal systems.

Together, these findings establish a molecular and functional framework for defining fibrocytes in cattle and demonstrate their suitability as a practical somatic cell source for reprogramming. By combining transcriptomic profiling with targeted culture optimization, this study reveals that fibrocytes represent an inducible hematopoietic lineage state with integrated immune, reparative, and metabolic programs that can be stabilized and expanded in vitro. The FbC medium maintains proliferative competence while suppressing inflammatory activation, providing a culture environment that preserves the intrinsic adaptability of fibrocytes and primes them for efficient induction of pluripotency. Beyond advancing reprogramming methodologies, these insights extend the understanding of fibrocyte biology, positioning them as a tractable interface between immune and stromal systems. Given their accessibility from peripheral blood and their stable propagation without genomic modification, bovine fibrocytes offer a scalable, field-adaptable platform for generating induced pluripotent stem cells across genetically diverse or valuable livestock. Future work defining the epigenetic state transitions that underlie fibrocyte plasticity and reprogramming will further clarify their role in regenerative and reproductive biotechnology and expand opportunities for genetic diversity preservation in cattle and other agriculturally important ruminants.

## Acknowledgements

We gratefully thank Geoffrey Hall, Manager of the Dairy Barn at Cornell University, for his expert assistance in blood draws from the cows used in these experiments.

## Author contributions

V.S., H.S., and P.P.K conceived the study and designed the experiments. H.S., S.G., and V.V.P. performed fibrocyte cultures from blood. H.S. performed phenotypic characterization, culture medium optimization and reprogramming experiments. P.P.K. performed the transcriptomics and data analyses. H.S., P.P.K., and V.S. analyzed the data and interpreted the results. H.S. and V.S. wrote the initial draft of the manuscript, and all authors contributed to editing and approved the final version. V.S. supervised the project.

## SUPPLEMENTARY INFORMATION

### Supplementary Tables

**Table S1.**
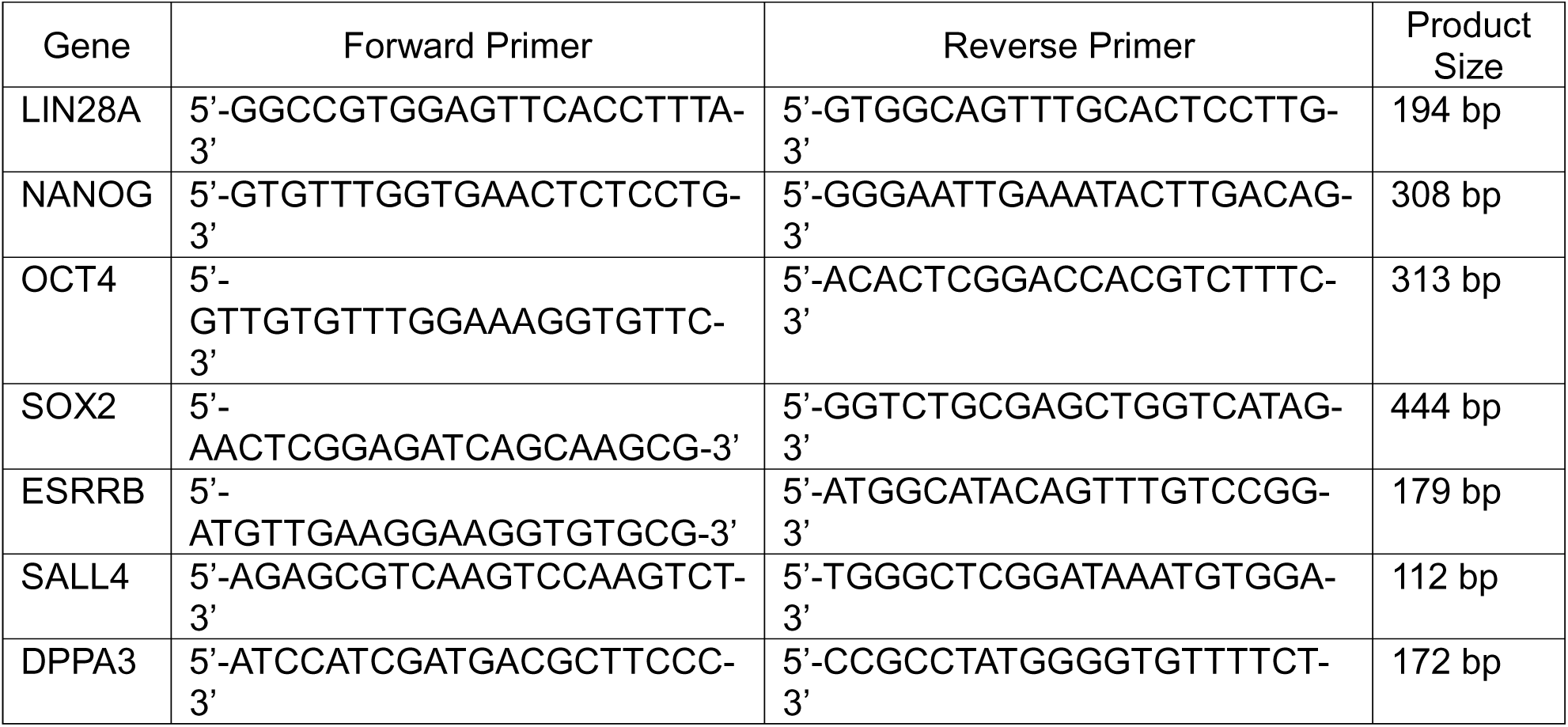
Primers used for RT-PCR.

### Supplementary Data File

**Data File S1. Transcriptomic distinctions between fibrocytes and fibroblasts. Tab 1:** Genes differentially expressed between fibrocytes (in Fb medium) and fibroblasts. **Tab 2:** Canonical pathways enriched in fibrocytes compared to fibroblasts. **Tab 3:** Prediction of upstream transcriptional regulation driving the gene expression pattern in fibrocytes. **Tab 4:** Prediction of upstream receptors driving the gene expression pattern in fibrocytes. **Tab 5:** Fibrocyte-specific gene expression data used to formulate functional gene networks.

**Data File S2. Transcriptomic effects driven by fibrocyte medium (FbC) conditions. Tab 1:** Genes differentially expressed between fibrocytes in FbC vs Fb medium conditions. **Tab 2:** Canonical pathways enriched in fibrocytes in FbC medium compared to fibrocytes in Fb medium. **Tab 3:** Genes regulated by FbC medium factors: dexamethasone, WNT signaling, TGFB1 inhibition, and EGF signaling.

### Supplementary Video Legends

**Movie S1. Fibrocyte outgrowths from peripheral blood-derived cells.** Time-lapse phase-contrast imaging of adherent peripheral blood-derived cells proliferating from a nucleation site on a gelatin-coated culture plate maintained in FbC medium. During the first 48 hours, numerous non-adherent cells and debris are present while attachment occurs. The sequence begins 24 hours after plating, showing the initial appearance of small adherent cell populations. A wash and medium change, indicated in the video, removes most non-adherent material. In this field, a nucleation site resembling a colony-forming unit becomes evident, from which fibrocytes proliferate outward and actively migrate across the plate, while the central region remains densely populated due to sustained proliferation. Images were acquired every 10 minutes and compiled at 7 frames per second, with the lower-right time stamp indicating elapsed minutes. Scale bar = 10 µm.

**Movie S2. Fibrocyte proliferation in Fb medium.** Time-lapse phase-contrast imaging showing fibrocyte proliferation in Fb medium. Fibrocytes previously maintained in Fb medium for approximately 10 days were passaged once and seeded at 3.0 x 10^4^ cells per well in a 24-well plate. The video captures cell migration, progressive attachment, elongation, and proliferation of fibrocyte-derived cells, with frames highlighting morphological transitions during cell division. The population shows a diffuse growth pattern with spindle-shaped cells displaying broadened lamellipodia and thick filopodia. Over time, extensive migratory activity and morphological changes becomes evident, with formation of polygonal cells and with some cells enriched in stress fibers. Images were acquired every 10 minutes for 7 days and compiled at 5 frames per second. Scale bar = 600 µm.

**Movie S3. Fibrocyte proliferation in FbC medium.** Time-lapse phase contrast imaging showing fibrocyte proliferation in FbC medium. Fibrocytes previously maintained in FbC medium for approximately 10 days were passaged once and seeded at 3.0 x 10^4^ cells per well in a 24-well plate. The video shows gradual cell attachment and spreading, with specific frames highlighting distinct mitotic events. The culture is composed of fusiform, squamous, and pyramidal-shaped cells exhibiting organized palisading projections. Compared to Fb medium, cells display reduced migratory behavior and enhanced proliferative activity. Images were acquired every 10 minutes over 7 days and compiled at 5 frames per second. Scale bar = 600 µm.

