## Supplementary material for "From blood to pluripotency: Fibrocytes as a reprogrammable somatic cell source for bovine iPSCs": Table S1

**Table S1. Primers used for RT-PCR**

| Gene | Forward Primer | Reverse Primer | Product Size |
| --- | --- | --- | --- |
| LIN28A | 5’-GGCCGTGGAGTTCACCTTTA-3’ | 5’-GTGGCAGTTTGCACTCCTTG-3’ | 194 bp |
| NANOG | 5’-GTGTTTGGTGAACTCTCCTG-3’ | 5’-GGGAATTGAAATACTTGACAG-3’ | 308 bp |
| OCT4 | 5’-GTTGTGTTTGGAAAGGTGTTC-3’ | 5’-ACACTCGGACCACGTCTTTC-3’ | 313 bp |
| SOX2 | 5’-AACTCGGAGATCAGCAAGCG-3’ | 5’-GGTCTGCGAGCTGGTCATAG-3’ | 444 bp |
| ESRRB | 5’-ATGTTGAAGGAAGGTGTGCG-3’ | 5’-ATGGCATACAGTTTGTCCGG-3’ | 179 bp |
| SALL4 | 5’-AGAGCGTCAAGTCCAAGTCT-3’ | 5’-TGGGCTCGGATAAATGTGGA-3’ | 112 bp |
| DPPA3 | 5’-ATCCATCGATGACGCTTCCC-3’ | 5’-CCGCCTATGGGGTGTTTTCT-3’ | 172 bp |
